# Airway injury induces alveolar epithelial responses mediated by macrophages

**DOI:** 10.1101/2024.04.02.587596

**Authors:** Irene G. Wong, Margherita Paschini, Jillian Stark, Alan Baez Vazquez, VanNashlee Ya, Aaron L. Moye, Susanna M. Dang, Maria F. Trovero, Emma L. Thompson, Sidrah Ahmed, Fatima N. Chaudhry, Andrea Shehaj, Maral J. Rouhani, Roderick Bronson, Sam M. Janes, Samuel P. Rowbotham, Jarod A. Zepp, Ruth A. Franklin, Carla F. Kim

## Abstract

Airway injury activates local progenitors and stimulates cell-cell interactions to restore homeostasis, but it is unknown how distal niches are impacted. We utilized mouse models of airway-specific epithelial injury to examine secondary tissue-wide alveolar and immune responses. Single-cell transcriptomics and *in vivo* validation of mouse models of airway-specific epithelial injury revealed transient, tissue-wide proliferation of alveolar type 2 (AT2) progenitor cells after club cell-specific injury or ablation. Myeloid cells exhibited altered gene expression after club cell loss and were detectable in the bronchoalveolar lavage fluid. The AT2 cell proliferative response was reliant on alveolar macrophages (AMs) exhibiting an injury-induced gene expression program. Overall, these results demonstrate that acute airway damage can trigger myeloid-mediated lung alveolar responses that may contribute to disease susceptibility or dysfunction.

## Introduction

In the lung and other tissues, injury and disease states promote cellular responses that occur locally to stimulate progenitor cells for repair of damaged epithelia, and in some scenarios, tissue-wide compensatory growth^1,2,3,4,5^. The lungs contain multiple epithelial cell types that act as stem/progenitor cells in the injured niche. It is not known if or how local epithelial injuries can affect changes in more distant epithelia. This may be particularly relevant for complex lung diseases like chronic obstructive pulmonary disease which exhibit dysfunction across multiple niches of the lungs, from the upper airways, small airways, to alveoli^6^. Club cells are epithelial stem cells that maintain the airway barrier^7^. Club cells can self-renew, differentiate into ciliated cells^8,9^ and in severe injury give rise to alveolar type 2 (AT2) cells^10^. Studies of airway injury and repair have focused on local responses around the airways^11^. The alveolar region contains AT2 epithelial cells that can self-renew and differentiate into alveolar type 1 (AT1) cells^7^. Whether and how AT2 cells are affected by airway injuries in the proximal or distal lung is unknown.

Club cells and AT2 cells are regulated within their respective niches by diverse cell types and factors, such as immune and mesenchymal cells. Monocytes are required for tracheal epithelial repair after polidocanol injury and tissue-wide proliferation after pneumonectomy^12,13^. After club cell-specific naphthalene airway injury, macrophage/monocytes increase in the lungs^14,15^ and depletion of alveolar macrophages (AMs) during the regenerative phase impaired airway repair^14,16^. Whether immune cells stimulated by airway epithelial defects specifically alter the alveolar space, however, is not known. Here we reveal immune cells as drivers of previously unknown alveolar responses to acute airway injury.

## Results

### Transient AT2 cell proliferation after naphthalene injury

To study tissue-wide effects of airway injury, we performed single cell RNA-seq on mouse lung epithelium from the naphthalene club cell injury model. We treated adult mice with corn oil diluent or naphthalene. Maximal loss of club cells occured 2 days post-naphthalene, detected by the absence of CCSP+ club cells in airways. Club cells regenerated over a period of up to 14-21 days (Figure 1A-B). We sorted alveolar-enriched Sca1- epithelium and airway-enriched Sca1+ epithelium^17^ during damage peak (day 2) and regenerative phases (day 7) for scRNA-seq (Figure 1A, S1A). Cell clusters were annotated with known markers for lung epithelial cell types^17^ (Figure S1B, Supplementary Table 1).

**Figure 1.**
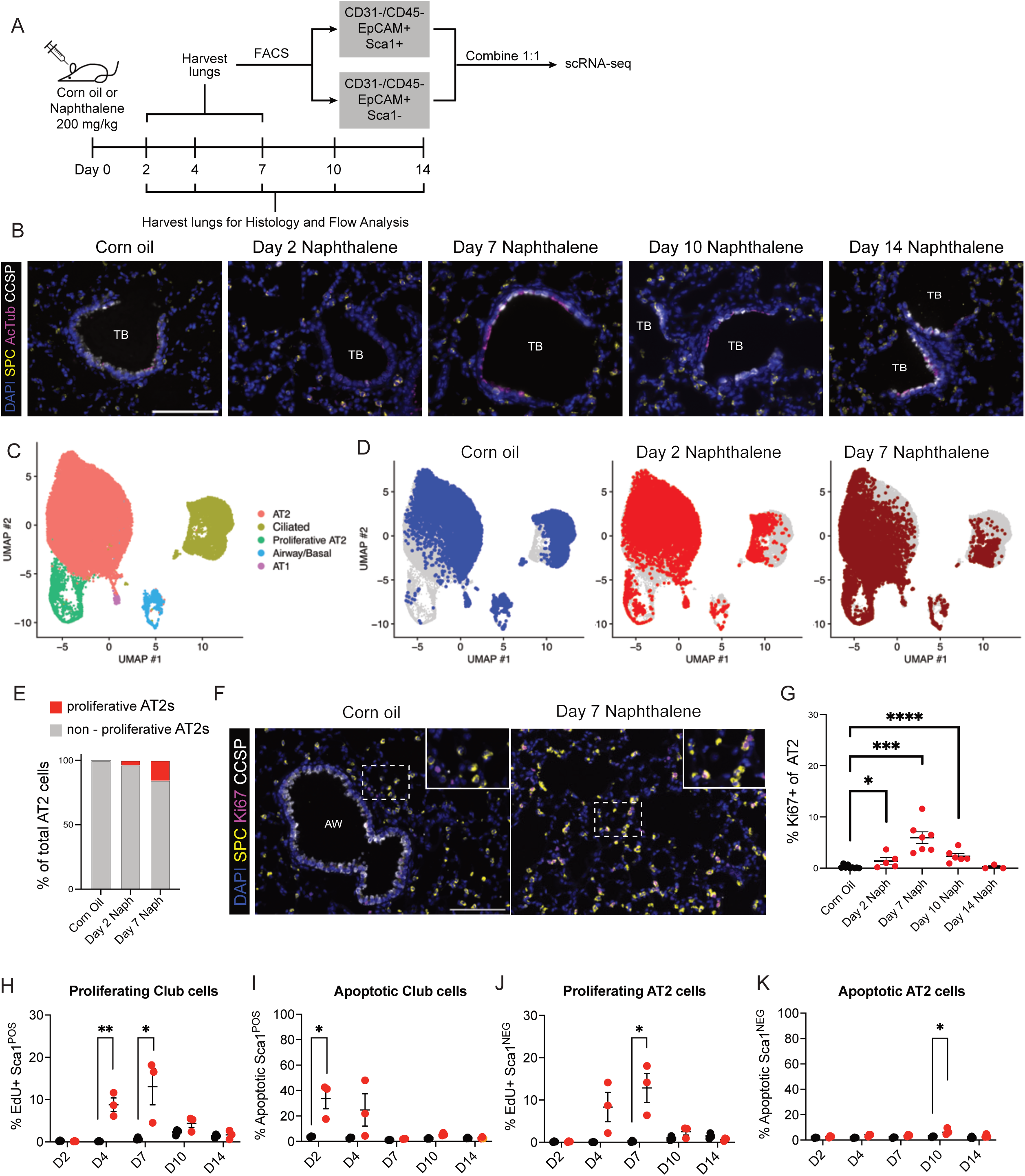
**Transient proliferation of AT2 cells after naphthalene airway injury.** A. Experimental design for Naphthalene injury experiments. B. IF images of Corn oil control or Naphthalene-treated mouse lungs. TB = Terminal bronchiole. Scale bar, 100 µm. C. Merged UMAP plot of epithelial cells from scRNAseq. n= 2 mice per condition. Corn oil N = 11216 cells; Day 2 Naphthalene N = 9411 cells; Day 7 Naphthalene N = 10145 cells. D. Split of the UMAP plot from C. based on condition: Corn Oil, Day 2 Naphthalene and Day 7 Naphthalene. E. Fraction of cells in the AT2 population of each condition determined by scRNA-seq data. F. IF images of Corn oil control mice and D7 Naphthalene-treated mice. Scale bar = 100 µm. G. Quantification of proliferative AT2 cells (Ki67+/SPC+) normalized to total AT2 cells at the indicated timepoints after naphthalene injury. Corn oil N = 9 mice, Day 2 N = 5 mice, Day 7 N = 7 mice, Day 10 N = 6 mice. Day 14 N = 3 mice. Student’s T-test. H. *In vivo* quantification of proliferating cells after Naphthalene injury by incorporation of EdU in the Sca1^POS^ population. N = 3 mice per condition, per timepoint. Student’s T-test. I. *In vivo* quantification of apoptotic cells after Naphthalene injury by AnnexinV staining in the Sca1^POS^ population. N = 3 mice per condition, per timepoint. Student’s T-test. J. *In vivo* quantification of proliferating cells after Naphthalene injury by incorporation of Edu in the Sca1^NEG^ population. N = 3 mice per condition, per timepoint. Student’s T-test. K. *In vivo* quantification of apoptotic cells after Naphthalene injury by AnnexinV staining in the Sca1^NEG^ population. N = 3 mice per condition, per timepoint. Student’s T-test.

We identified epithelial populations including club/goblet cells, AT2 cells, BASCs, PNECs and basal cells (Figure 1C, S1C, Supplementary Table 2), with changes reflective of naphthalene injury and subsequent responses. A cluster characterized by the expression of Scgb1a1 (Secretory Cells I) was largely constituted of cells from controls (Figure S1D), reflecting homeostatic airway. This cluster was depleted upon injury, with few cells captured on day 2. On day 7, secretory cells populated another cluster (Secretory Cells III), likely newly generated club cells (Figure S1D). A third secretory cluster (Secretory Cells II) expressed a mucin/goblet cell signature and was not affected by naphthalene. Surprisingly, proliferative AT2 cells were identified after airway injury (Figure 1D-E, S1E). Naphthalene also stimulated an AT2 primed cell state, previously associated with injury,^18–20^ that further increased 7 days post naphthalene (Figure S1F). There was no evidence for cells in the alveolar transitional/intermediate state^18–20^ (Figure S1F). Because naphthalene is only metabolized by club cells that express Cyp2f2 and converted into toxic reactive oxidative species^21,22^, changes to AT2 cells were not expected. Cyp2f2 was detected only in CCSP+ club cells, suggesting that AT2 cells do not metabolize naphthalene (Figure S1G).

Immunofluorescence (IF) confirmed observations from the scRNA-seq, showing that AT2 cell proliferation occurred after naphthalene-mediated club cell injury. IF for Ki67^23^ and the AT2 cell marker SPC was performed at various time points. The percentage of AT2 cells in active proliferation (SPC+ Ki67+) increased at day 2 post injury, reached maximum value at day 7 and eventually subsided by day 14 (Figure 1F-G). We used CCSP-driven Cre recombinase to lineage trace club cells prior to naphthalene injury^24^. We did not observe lineage-labeled YFP+ club cells in the alveolar space at day 7 after injury (Figure S1H-J); AT2 cell proliferation after naphthalene-mediated club cell injury occurred in native AT2 cells and was not from differentiated club cells.

To orthogonally examine the impact of airway injury on AT2 cells, we performed *in vivo* labeling of proliferating cells and detection by flow cytometry. Mice were treated with corn oil or naphthalene and injected with EdU at 12 and 24 hours before harvest to label cells with active DNA synthesis (Figure 1A). Populations enriched for AT2 or club cells (DAPI- CD31/45- EpCam+ Sca1- for AT2; Sca1+ for Club cells) were assessed for EdU+ cells (Figure S1A) and cell death by AnnexinV staining. As expected, club cells had a large portion of apoptotic cells 2-4 days after injury, closely followed by a significant burst of proliferation starting at day 4 (Figure 1H-I). These findings are consistent with the initial acute damage and subsequent repair response that the airways are known to undergo after naphthalene injury. Apoptosis of AT2 cells was not detected at early time points after naphthalene injury. AT2 cells exhibited a substantial proliferative response between days 4 and 7, as shown by the significant uptick in EdU-labeled cells, confirming the sequencing and IF observations (Figure 1J-K).

### Transient AT2 cell proliferation after genetic club cell ablation

We next used a genetic model of club cell ablation to corroborate our findings regarding AT2 cell proliferation in the chemical model. In CCSP-CreER+/-; Rosa26-LSL-YFP/iDTR mice^24,25^, tamoxifen induced expression of the diphtheria toxin receptor (iDTR) and YFP label on CCSP-expressing club cells (Figure S1G). We administered PBS (control) or diphtheria toxin-A (DT) to acutely deplete club cells in these mice (Figure 2A). Lineage labeling, also a proxy for iDTR, was confined to club cells and did not label any SPC+ AT2 cells (Figure S2A). CCSP+ club cell injury peaked at day 2 after DT administration, with no other obvious histological changes in the injured lungs compared to controls (Figure 2B-C, Figure S2B-C). AT2 cell proliferation peaked at day 4 after club cell ablation (Figure 2D-E). The increase in AT2 cell proliferation after club cell ablation was transient, disappearing as airways underwent repair. Despite differences in the mechanism and severity of club cell injury between the naphthalene and genetic ablation, these results demonstrated AT2 proliferation in response to club cell depletion in two different models.

**Figure 2:**
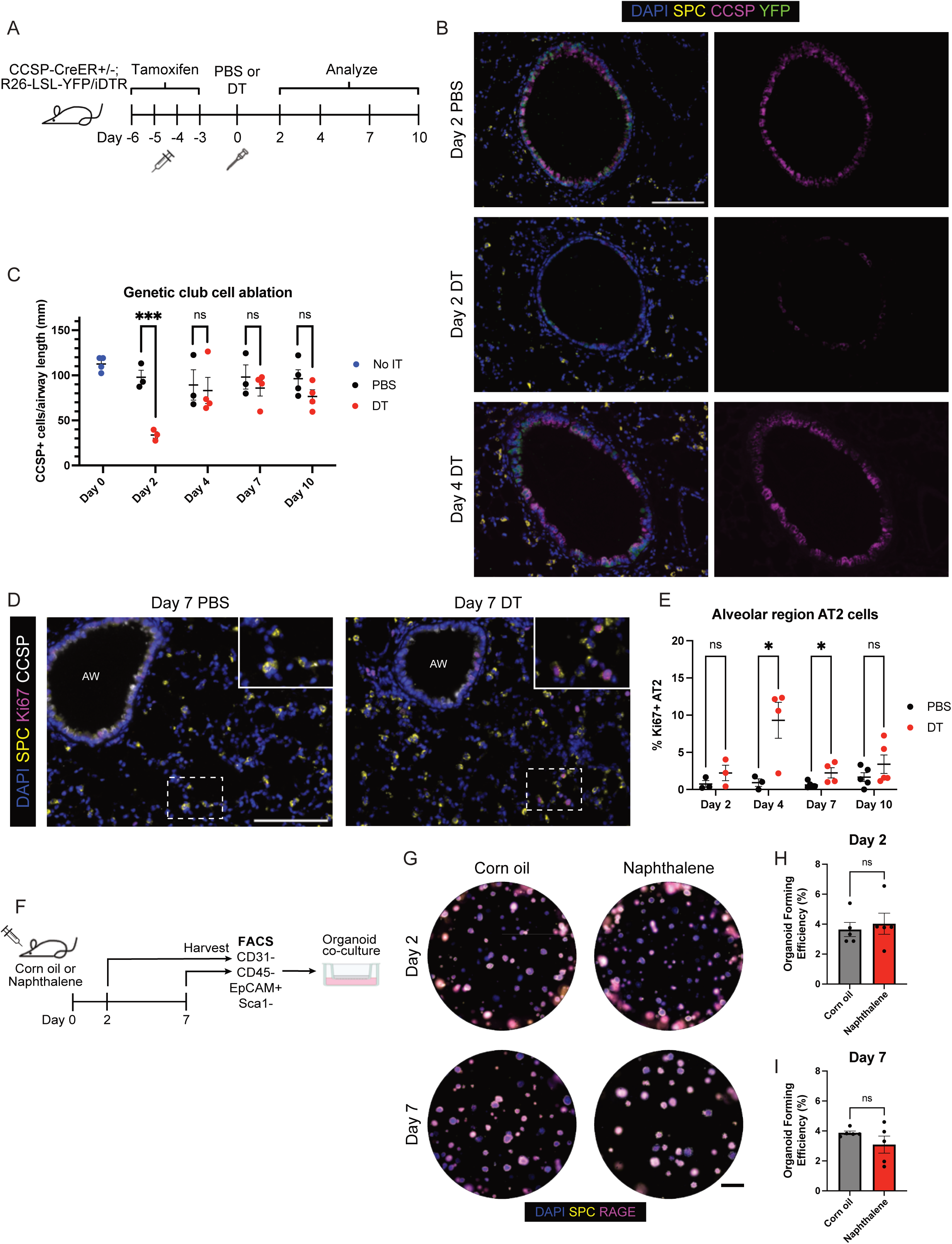
**Transient proliferation of AT2 cells after genetic club cell ablation.** A. Experimental design of induction and injury of CCSP-CreER+/-; Rosa26-LSL-YFP/iDTR mice. Tamoxifen was administered via intraperitoneal injection. PBS or DT were administered via intratracheal injection. B. IF images of mouse lungs at the indicated timepoints after genetic club cell ablation. Scale bar = 100 µm. C. Quantification of CCSP+ club cell loss in airways of CCSP-CreER+/-; Rosa26-LSL-YFP/iDTR mice after no intratracheal administration, PBS, or DT administration at the indicated timepoints. N = 3-4 mice per condition, per timepoint. Student’s T-test. ns = no significance. D. IF images of PBS control mice and DT-treated mice sacrificed at day 7 after injury. Scale bar = 100 µm. E. Quantification of proliferative AT2 cells (Ki67+/SPC+) normalized to total AT2 cells at the indicated timepoints after genetic club cell injury. N = 3-5 mice per condition, per timepoint. Student’s T-test. ns = no significance. F. Experimental design of isolating alveolar-enriched Sca1- lung epithelial cells from corn oil control or naphthalene-treated mice and setting up 3D organoid co-culture with stromal cells. G. IF of alveolar 3D organoid co-cultures grown from Sca1- lung epithelium as diagrammed in (F). Scale bar = 1 mm. H. Quantification of organoid forming efficiency of Sca1- alveolar organoids collected at day 2 after Naphthalene injury as shown in (G). N = 5 mice per condition, per timepoint. Student’s T-test. ns = no significance. I. Quantification of organoid forming efficiency of Sca1- alveolar organoids collected at day 7 after Naphthalene injury as shown in (G). N = 5 mice per condition, per timepoint. Student’s T-test. ns = no significance.

To test whether AT2 cell proliferation was a cell-autonomous behavior, we isolated alveolar-enriched epithelium from control or naphthalene-treated mice on day 2 or 7 and tested their ability to form organoids^26^ (Figure 2F). Organoid forming efficiency (OFE) was unchanged whether the epithelial cells were derived from injured or uninjured animals, suggesting additional cues from the injured environment are required to promote increased AT2 cell proliferation (Figure 2G-I).

### Multiple myeloid lineages respond to airway injury

We sought to interrogate potential contributions of immune populations to the AT2 cell proliferation after club cell injury by single cell transcriptomic analysis. Previous work demonstrated the importance of immune cells in promoting lung epithelial cell proliferation in other injury models^16,27–29^. To capture immune cells from the airspace and within lung tissue, scRNA-seq was performed on a mixture of cells from bronchoalveolar lavage fluid (BALF) and digested lung from control, days 2 and 7 after naphthalene (Figure 3A, Figure S3A). We identified major immune cell types leveraging published datasets^30,31^ (Figure 3B, Figure S3B-C, Supplementary Table 3). Lymphoid populations from injured mice, such as T cells, B cells, Natural Killer cells and type 2 innate lymphoid cells, did not show significant changes when compared to those from control mice (Figure 3D). In contrast, myeloid immune cells from injured mice, including macrophages and neutrophils, exhibited significant gene expression changes.

**Figure 3:**
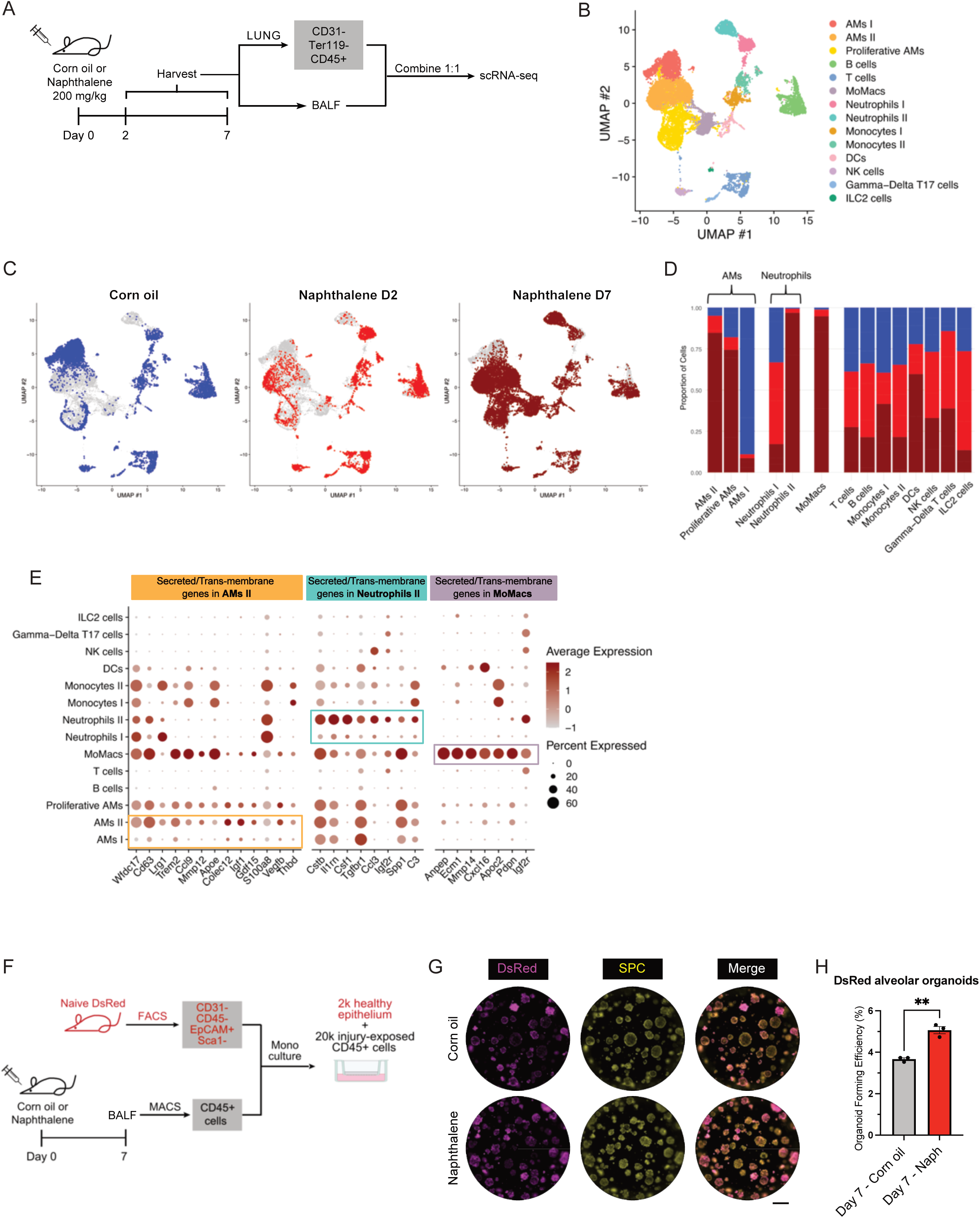
**Multiple myeloid lineages respond to airway injury.** A. Experimental design for isolating AMs (via BALF) and remaining CD45+ immune cells in the lungs from mice treated with corn oil control or naphthalene for scRNA-seq. B. Merged UMAP plot of immune cells from (A). N = 3 mice per condition. Corn oil N = 6626 cells; Day 2 naphthalene N = 4185 cells; Day 7 Naphthalene N = 11200 cells. C. Split of the UMAP plot from (B) based on condition. D. Composition of each cluster by condition. E. Dot Plot of secreted and transmembrane genes strongly enriched in the AMs II (left panel), Neutrophils II (central panel) and MoMacs clusters (right panel). F. Experimental design for isolating immune cells from BALF using CD45-positive selection with MACS. These cells were combined with Sca1- alveolar-enriched epithelium from naïve DsRed animals to establish 3D organoids co-cultures. G. IF of alveolar 3D organoid co-cultures grown from Sca1- lung epithelium as diagrammed in (F). Scale bar = 1 mm. H. Quantification of organoid forming efficiency of alveolar organoids shown in (G). Student’s T-test.

AMs were distributed into three clusters based on their distinct gene expression program and cell cycle status induced by naphthalene injury. Cluster AM I cells were mostly from control corn oil-treated mice, cluster AM II was composed mostly of AMs from day 7, and proliferative AMs, highly expressing genes associated with active cell cycling, were nearly all from injury samples (Figure 3C-D, S3C). To identify signaling molecules from AMs that may contribute to AT2 proliferation after naphthalene, a list of genes upregulated in the AM I cluster vs the AM II cluster (Supplementary Table 4) was further refined for secreted or transmembrane proteins (Figure 3E, left). Multiple secreted proteins (Lrg1, Wfdc17, Apoe, S100a8, Vegfb), cytokines (Gdf15 and Ccl9), elastin degrader Mmp12 and transmembrane proteins (Cd63, Thbd, Trem2 and Colec12) were upregulated. Many of these candidates were also upregulated in proliferative AMs. IL-1B (*Il1b),* implicated in alveolar repair^18^, was only minimally enriched in AMs after naphthalene-mediated injury (Figure S3D).

Neutrophils also underwent a significant transcriptional shift after naphthalene. This was especially evident at day 7, when a cluster of neutrophils was identified that were not present in control or day 2 samples (Figure 3B-3D, Figure S3B). Genes that differentiated injury-specific neutrophils from controls (Neutrophils I vs Neutrophils II, Supplementary Table 5) were filtered for secreted or transmembrane proteins (Figure 3E, middle). Multiple cytokines and chemokines dominated the list of naphthalene-specific genes (Cstb, Csf1, Il1rn, Ccl3, C3 and Spp1), suggesting orchestration of a myeloid inflammatory program.

Finally, we noted the emergence of a cluster of cells reminiscent of Monocyte-derived Macrophages (MoMacs) coincident with the emergence of proliferative AT2 cells. MoMacs are recruited to the airspaces upon alveolar injury and are capable of maturing into AMs^32^. Cells with the gene expression program of MoMacs appeared 7 days after airway damage (Figure 3C-D). Potential signaling nodes active in MoMacs included secreted factors (Ecm1, Apoc2, Cxcl16), along with known secreted ECM-remodeling enzymes (Mmp14) and transmembrane proteins (Pdpn, Igfr2 and Anpep) (Figure 3E, right panel). Interstitial macrophages were not resolved as a distinct cluster.

### Myeloid cells reprogrammed by airway injury impact AT2 cell proliferation

Sequencing analysis showed myeloid cells exhibited the most significant changes after naphthalene, therefore we examined their relative abundance using flow cytometry. Within CD45+ cells, AMs were identified by Ly6g- CD64+ MertK+ SiglecF+ Cd11b-, Neutrophils by Ly6g+, and IMs and MoMacs by Ly6g- CD64+ MerTK+ SiglecF- Cd11b+ (Figure S3E). We observed a significant increase in the percentage of the CD45-positive fraction detected as AMs at day 4 and a decreased fraction of AMs at day 10, coincident with the timing that preceded and followed the AT2 proliferative wave (Figure S3F). The percentage of IMs/MoMacs was also increased at day 4. In the genetic ablation model, the percentage of AMs and IMs/MoMacs did not change, yet neutrophils were significantly increased (Figure S3G).

We performed functional assays to determine the impact of myeloid cells on AT2 cells, by testing if immune cells isolated from BALF affected AT2 organoid formation^33^. BALF CD45+ cells from control or naphthalene-treated mice at day 7 after injury were enriched and used for co-culture with naïve DsRed+ AT2 cells in a growth factor-defined media^34^ (Figure 3F). BALF CD45+ cells from naphthalene-injured mice promoted a significantly higher AT2 cell OFE compared to BALF CD45+ cells from uninjured mice (Figure 3G-H). These results suggested that immune cells producing naphthalene-responsive factors can directly impart pro-proliferative effects on AT2 cells.

To further define components that mediate alveolar changes, we analyzed the cellular content of the BALF by flow cytometry. BALF was collected 7 days after naphthalene injury, when gene expression patterns were most significantly changed, to quantify the major myeloid populations. Control BALF was predominantly composed of AMs (∼80%), whereas BALF from naphthalene-treated mice contained neutrophils, MoMacs, and other un-identified immune cells (Figure 4A-B), as observed in other contexts of insult.^35,36^ Notably similar numbers of AMs were present in BALF from control and injured mice (Figure S4A). We also collected BALF from mice 4 days post DTA instillation when the AT2 proliferation peaks after genetic club cell ablation. We observed altered percentages of AMs and neutrophil infiltration without MoMacs (Figure 4C).

**Figure 4:**
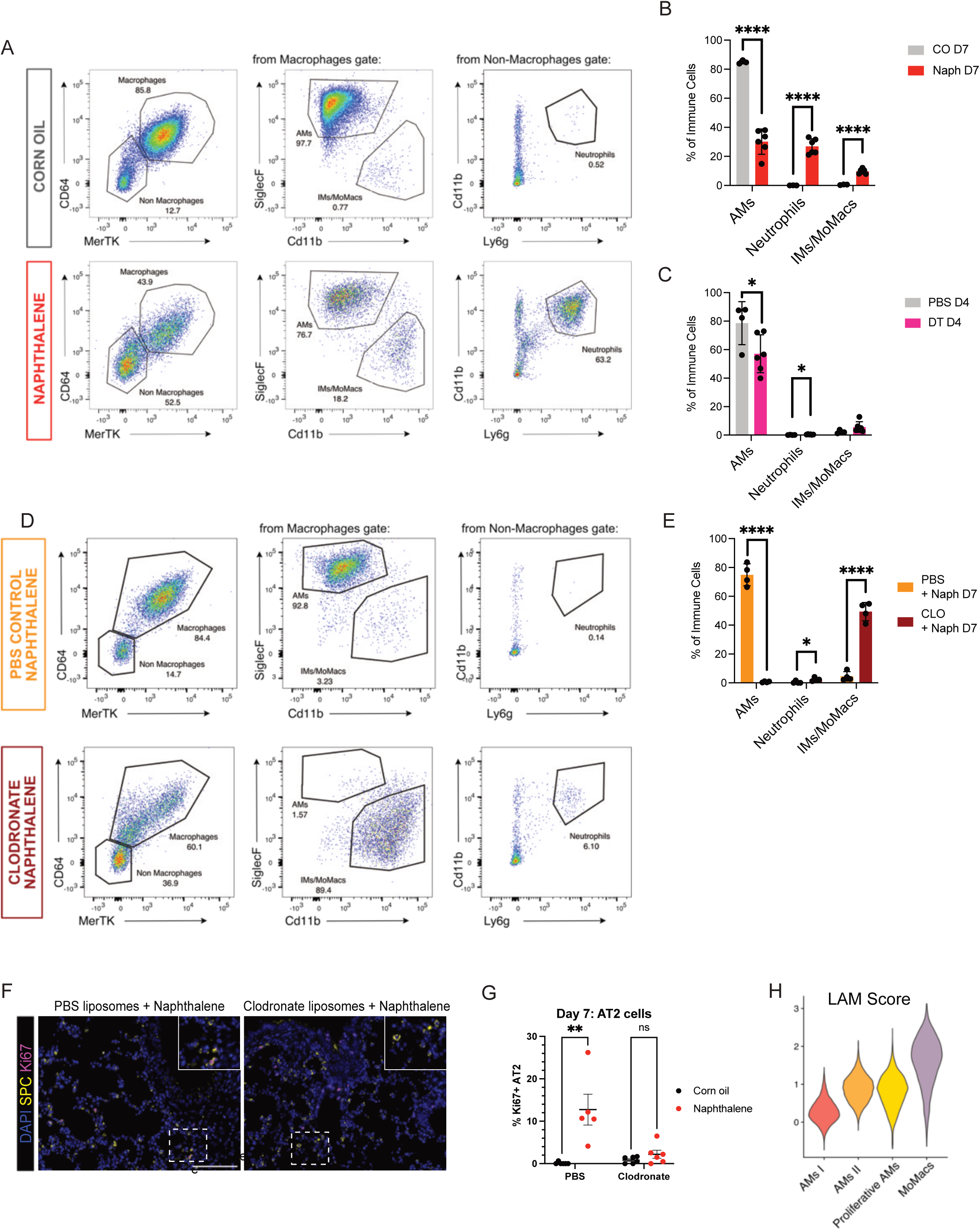
**Myeloid cells reprogrammed by airway injury impact AT2 cell proliferation.** A. Flow analysis and gating strategy of the myeloid population in BALF fluid from Corn Oil and Naphthalene mice 7 days post injury. Representative images. B. Quantification of the myeloid populations in BALF fluid of Corn Oil and Naphthalene treated mice 7 days post injury. N ≥ 3 mice per condition. Student’s T-test. C. Quantification of the myeloid populations in BALF fluid of PBS and DTA treated CCSP-CreER+/-; Rosa26-LSL-YFP/iDTR mice, 4 days post injury. N ≥ 3 mice per condition. Student’s T-test. D. Flow analysis of myeloid populations 7 days after Naphthalene treatment in mice pre-treated with PBS or Clodronate liposomes as described in S4B. E. Quantification of the myeloid populations in BALF fluid of mice treated with Naphthalene (D7 after injury) and pre-treated with PBS or Clodronate liposomes. N ≥ 3 mice per condition. Student’s T-test. F. IF images of mice treated with Naphthalene (D7 after injury) and pre-treated with PBS or Clodronate liposomes. Scale bar = 100 µm G. Quantification of proliferative AT2 cells (Ki67+/SPC+) normalized to total AT2 cells at day 7 (D7) after liposome pre-treatment and naphthalene injury. N = 5-6 mice per condition. Student’s T-test: ns = no significance. H. LAM signature score across the Macrophages clusters from the single cell analysis shown in Figure 3.

Given the ability of BALF, containing AMs, to affect AT2 cells in organoid cultures and the known critical role of AMs after repair in multiple lung injury models, we asked if AMs are required for the airway injury-induced AT2 cell response. We pre-treated mice with PBS control liposomes or clodronate liposomes to deplete macrophages from the lung airspaces prior to naphthalene injury and tested the effects on AT2 cell proliferation^14,37,38^ (Figure S4B). Similar club cell loss was seen in PBS and clodronate treated mice at day 7 after injury (Figure S4C). Pre-treated and injured lungs did not highlight obvious differences in lung morphology and structure (Figure S4D). To confirm the loss of AMs upon clodronate treatment, we collected BALF from mice at day 7 post-injury. In mice treated with PBS liposomes and naphthalene, BALF was largely composed of AMs, whereas in mice treated with Clodronate liposomes and naphthalene, BALF almost completely lacked AMs (Figure 4D-E, S4E). Clodronate liposome treatment attenuated AT2 proliferation responses to naphthalene at day 7 after injury (Figure 4F-G). Notably, with clondronate liposome treatment and the ablation of AMs, a substantial increase in Monocyte-derived Macrophages (Ly6g- CD64+ MertK+ SiglecF- Cd11b^low^ cells) was seen (Figure S4E). Nevertheless, the ablation of AMs correlated with the absence of the AT2 proliferation, implicating AMs in the alveolar response to club cell loss.

To further understand the role of AMs in the airway injury response and in AT2 proliferation, we compared the transcriptional profile of AMs from control and naphthalene-treated mice. Pathway-level profiling using Reactome^39^ revealed a significant enrichment for lipid handling with terms “Cholesterol Biosynthesis” and “Metabolism of Steroids” as the most significant enriched pathways (Figure S4F). Lipid-associated macrophages (LAMs) identified in the lung after lipopolysaccharide, bleomycin injury and cigarette smoke, have been proposed to play a critical role after insults specific to the alveolar space^36,40–42^. We generated a signature for lung LAMs^40,43^ and scored the expression of this signature in the scRNA-seq macrophage clusters (Figure 3). AM II cluster and Proliferative AMs cluster exhibited an increased score for the LAM signature compared to the AMs I cluster, suggesting that upon naphthalene, AMs assume LAM characteristics (Figure 4H). Furthermore, MoMacs showed the highest enrichment in the LAM signature, consistent with literature reporting that LAMs emerge from monocyte-derived Macrophages. Since Spp1 was highly expressed in LAM-like macrophages after naphthalene, we used Spp1-CreERT2; LSL-Tom mice to understand the location of these cells^44^. We observed a robust presence of labeled cells in the alveolar space 7 days after Naphthalene, consistent with their putative role in regulating AT2 cells (Figure S4G).

## Discussion

In this study, we discovered depletion of airway epithelial cells leads to a complex lung alveolar response, including alveolar progenitor cell proliferation mediated by macrophages. In response to chemical or genetic loss of club cells, a variety of myeloid populations infiltrated the airspaces and acquired a gene expression program that included production of a variety of secretory factors. This injury-associated response imparts a temporary proliferative effect on AT2 cells distinct from cell states associated with alveolar differentiation. Clodronate experiments suggested AMs are necessary for regulating AT2 cell turnover. Similarly, a decrease in overall lung epithelial proliferation was seen upon AM depletion prior to naphthalene injury^16^. Multiple myeloid cell types, including neutrophils and MoMacs, are also altered following airway epithelial injury^14,16^.

AMs and other myeloid populations appear to use precise mechanisms to regulate progenitor cells, tailored to the injury context. Macrophages promote airway epithelial proliferation at the site of club cell injury for repair via signaling via an IL-33-ST2 axis, with IL-33 expression increased in club cells after naphthalene^14^. AMs can induce alveolar epithelial proliferation in response to lung alveolar damage after bleomycin injury, hookworm infection, pneumonectomy, and influenza^12,28,45^, with recent studies revealing AM-derived oncostatin M as a key driver of AT2 proliferation^27^. The effects of myeloid cells on AT2 cells downstream of airway injury appear to be distinct from those that stimulate the alveolar transitional cell state and AT1 cell differentiation and may be regulated by distinct mechanisms. We hypothesize similar proliferative effects over time could contribute to the limited abundance of alveolar progenitor cells and increased disease susceptibility known to be associated with age^46^. Consistent with this idea, naphthalene treatment prior to oncogenic Kras induction increased lung adenocarcinoma tumor burden and severity^47^. Further, injury or infections affecting the airway are known to precede the onset of lung diseases that impact the alveolar epithelia such as COPD and interstitial lung disease^48^.

In future studies, delineation of cocktails of factors that can influence specific AT2 cell responses such as proliferation or differentiation may be useful in regenerative medicine and disease treatment. We hypothesize AT2 cell proliferation after naphthalene is mediated by a combination of the secreted factors from the AMs in the LAM state, produced during the injury response. Among the LAM-associated factors, Igf1 was one of the most significantly enriched secreted genes in AMs from naphthalene-treated mice and has been shown to directly increase proliferation of AT2 cells in vitro and play important roles in the lung during development and aging^49–51^. Our work further underscores how the return to tissue homeostasis after an injury is a highly complex process, which affects epithelial cells and their interactions beyond the site of damage. Building upon this knowledge with mechanistic studies will illuminate how lung diseases develop and how to design improved therapies for patients.

## Limitations of the study

Pretreatment with clodronate significantly blunted the AT2 proliferative response after naphthalene airway injury. Clodronate depleted AMs, and we concluded that AMs are likely the conduit between the airway injury and the proliferation of the AT2 cells. However, we also noticed a significant infiltration of MoMacs with clodronate treatment. We cannot exclude a potential role of MoMacs in blocking the proliferation of AT2 cells or an important role of neutrophils. Experiments to block MoMacs recruitment with *Ccr2* knockout mice or antibody-mediated ablation of neutrophils would further define the role of myeloid populations in this phenomenon. We tested the role of macrophage-derived Spp1 using a conditional knockout strategy and found that AT2 cells were still proliferative; we have not yet determined which factor(s) are required for AT2 proliferation after airway injury or the molecular crosstalk from proliferating AT2s that may reprogram macrophages. We did not analyze immune cell changes at multiple time points in the genetic ablation model due to breeding challenges. Future studies could elucidate how the airway-alveolar-macrophage responses change with more severe, durable, or complex lung injuries and disease states.

## Acknowledgements

We thank Kim Lab members, Rania Dagher, Michal Shoskes-Carmel, Ramesh Shivdasani, Allon Klein, David Breault and Ke Yuan for thoughtful discussions; Ronald Mathieu, Mahnaz Paktinat, and Ranjan Maskey of the BCH Flow Cytometry Core; the Rodent Histopathology Core at Harvard Medical School; Kevin Bi and Preetida Bhetariya for data analysis advice; Frederic Lemaigre for Spp1-CreERT2 mice.

## Author Contributions

I.G.W. and M.P. designed, conceived, and performed experiments and wrote the manuscript. J.S., V.Y., A.B.V., S.M.D., M.J.R., M.F.T., S.A., E.L.T., F.N.C designed and performed experiments, M.P., I.G.W. and A.L.M. generated and analyzed scRNA-seq data. R.B. performed histopathological analysis. S.M.J., S.P.R., J.A.Z., R.A.F. and C.F.K supervised the design and study and co-wrote the manuscript. All authors reviewed and edited the manuscript.

## Resource Availability

### Lead Contact

The lead contact for this study is Dr. Carla F. Kim, PhD

### Materials Availability

### No unique material was generated in this study. <u>Data and Code Availability</u>

All raw and processed scRNA-seq data were deposited to the NCBI Gene Expression Omnibus (GEO) and Sequencing Read Archive (SRA) under the accession code GSE261506 and GSE261507. Bioinformatics analyses are available on GitHub https://github.com/mimapaschini/Naph_code.

## STAR Methods

### KEY RESOURCES TABLE

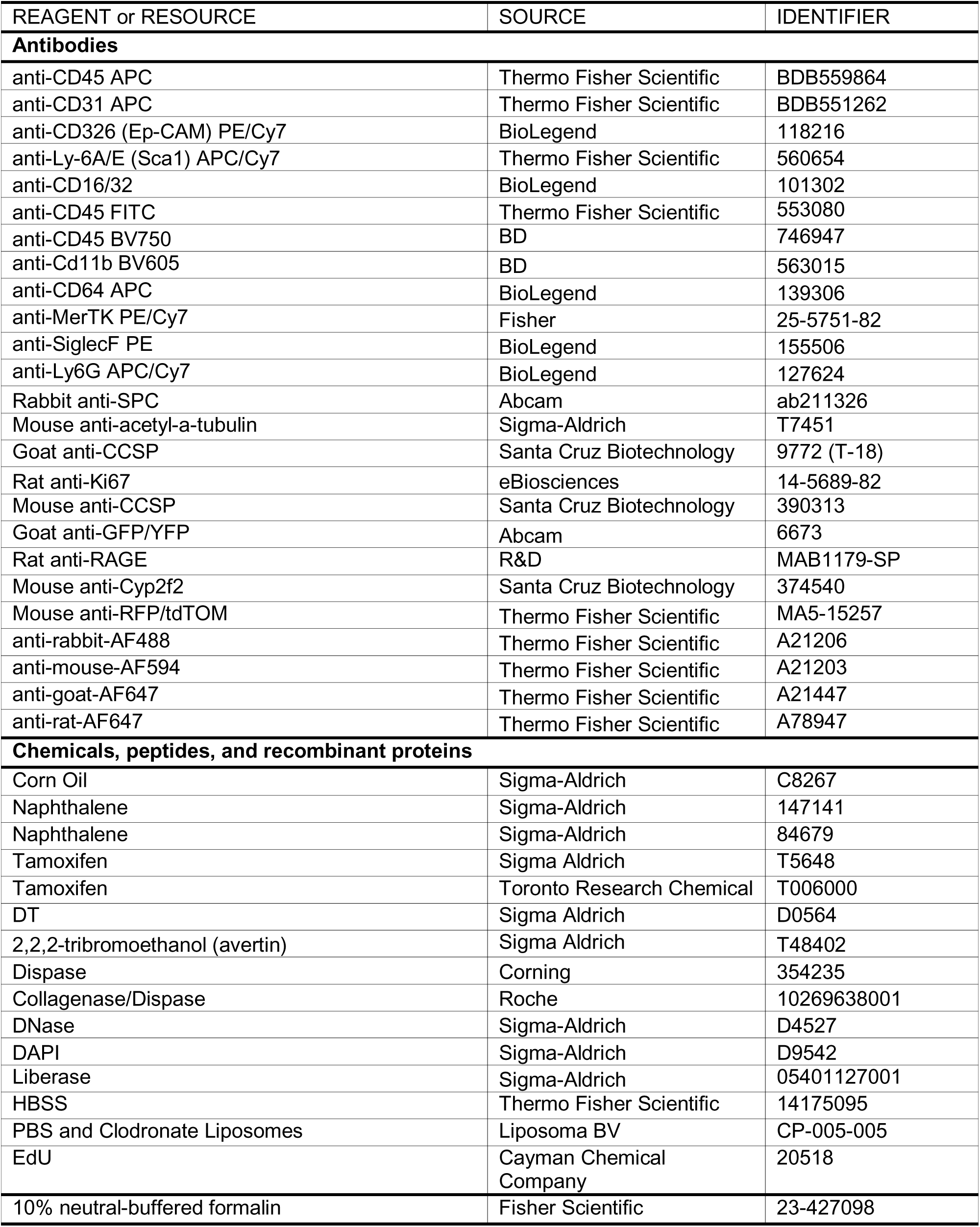
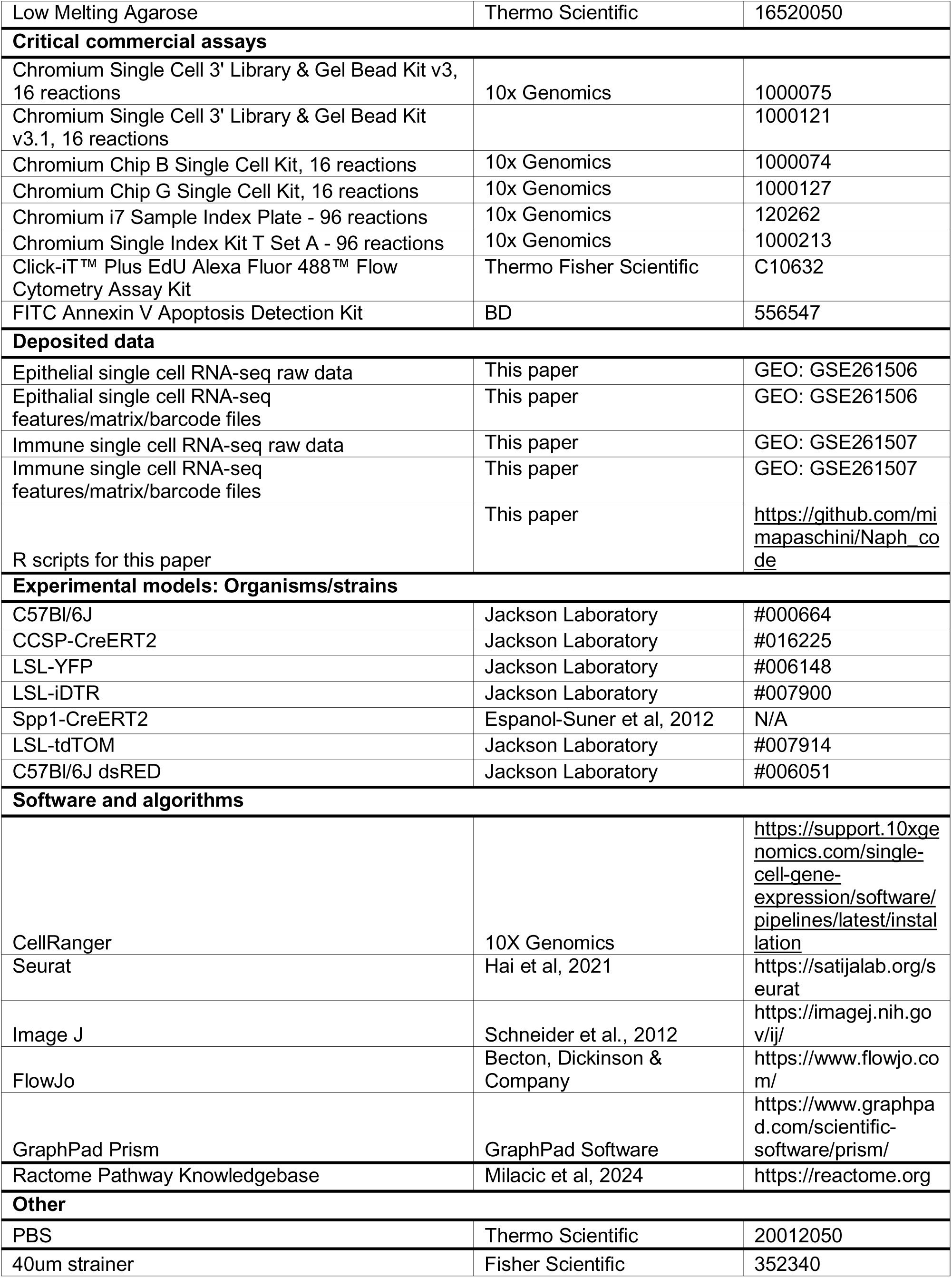
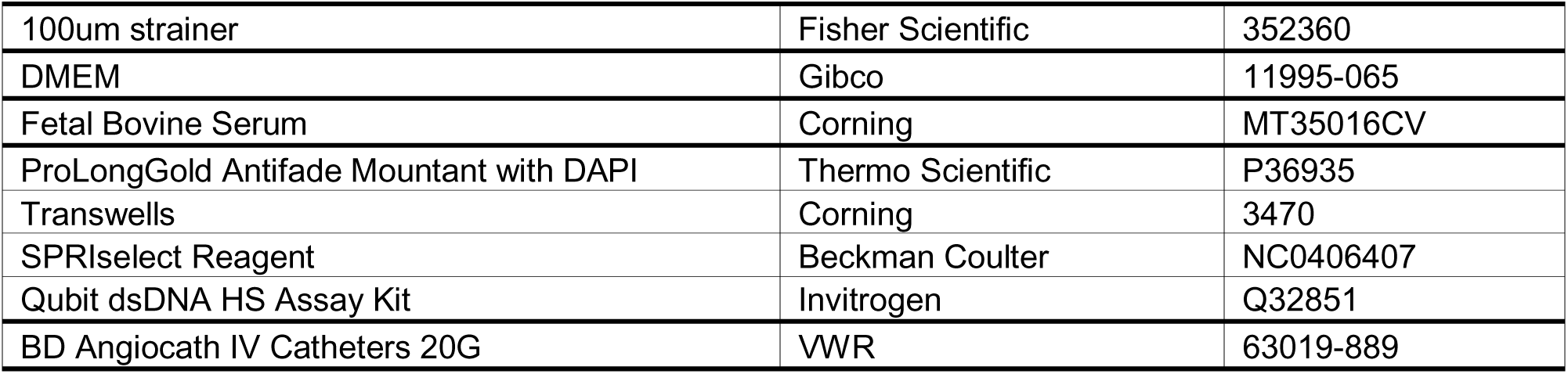

#### Experimental model and study participant details

##### Animals

Adult mice (8-12 weeks) were used for all studies. C57Bl/6J mice (Jackson Laboratory #000664) were utilized unless otherwise noted. CCSP-CreER (Jackson Laboratory #016225) mice were crossed to R26-LSL-YFP (Jackson Laboratory #006148) and maintained at homozygosity for both genes. CCSP-CreER; R26-LSL-YFP were then crossed to homozygous R26-LSL-iDTR (Jackson Laboratory #007900) mice to create CCSP-CreER+/-; R26-LSL-YFP/iDTR mice for genetic club cell ablation. SPP1-CreERT2 mice were generously donated by Frederic Lemaigre (Université Catholique de Louvain, Brussels, Belgium) and crossed to homozygous R26-LSL-tdTOM (Jackson Laboratory #007914). C57Bl/6J dsRED mice were used to obtain dsRED AT2 cells (Jackson Laboratory #006051). All mice were maintained in virus-free conditions and housed in groups. All experiments were approved by the Boston Children’s Hospital and Children’s Hospital of Pennsylvenia Animal Care and Use Committee. Due to documented sex differences in the response to naphthalene-mediated club cell injury, female mice were utilized for injury studies and grouped by litter when possible.

#### Method details

##### Naphthalene Injury

Naphthalene (Sigma Aldrich #84679) was dissolved in corn oil (Sigma Aldrich #C8267) at a concentration of 20 mg/mL at room temperature and sterile filtered before administration. C57Bl/6J or CCSP-CreER+/-; R26-LSL-YFP/iDTR mice were injected intraperitoneally with naphthalene at 200 mg/kg body weight or an equivalent volume of corn oil for control animals. Mice were then monitored for signs of distress throughout recovery until sacrificed with anesthetic overdose at indicated timepoints.

##### Genetic club cell ablation

CCSP-CreER+/-; R26-LSL-YFP/iDTR were first induced by 4 intraperitoneal injections of tamoxifen (Sigma Aldrich T5648) administered once daily, at 100 mg/kg body weight (stock solution 20 mg/mL in corn oil). On the third day after the last dose of tamoxifen, diphtheria toxin (Sigma Aldrich D0564) was administered via intratracheal injection at 3 µg/kg body weight (stock solution 60 µg/mL in PBS) or PBS control (Thermo Fisher #20012050). Mice were then monitored for signs of distress throughout recovery and sacrificed at indicated timepoints.

##### In vivo labeling of proliferating cells using EdU

For detection of proliferating cells, EdU (1.5 mg/dose in 200ul of PBS, Cayman Chemical Company #20518) was given intraperitoneally 24 and 12 hours before harvesting. For detection of proliferating cells via incorporation of EdU in vivo, the Click-iT™ Plus EdU Alexa Fluor 488™ Flow Cytometry Assay Kit (Thermo Fisher Scientific C10632) was used according to the manufacturer’s protocol. After standard Epithelial staining, cells were fixed and permeabilized. Incorporated EdU was detected via a copper-catalyzed azide-alkyne cycloaddition (Click-iT™ chemistry) with an Alexa Fluor 488™ azide, allowing direct fluorescent visualization of proliferating cells.

##### In vivo labeling of Spp1-expressing cells

Spp1-CreERT2; LSL-tdTOM mice were injected intraperitoneally with naphthalene at 200 mg/kg body weight (Sigma Aldrich 147141) or an equivalent volume of corn oil for control animals. To label Spp1-expressing cells, each mouse was induced by oral gavage of tamoxifen (Toronto Research Chemical #T00600) administered at -1, 0, +1 and +4 days from injury with at 200 mg/kg body weight (stock solution 20 mg/mL in corn oil).

##### Lung tissue preparation, flow cytometry, FACS for Epithelial cells

Lungs were dissected, prepared, stained, analyzed, and sorted as described previously^17,26^. Briefly, mice were anesthetized with avertin overdose (Sigma Aldrich #T48402). Lungs were perfused with cold PBS followed by intratracheal instillation of 2 mL dispase (Corning #354235). Lungs were placed on ice, minced, and incubated in 0.0025% DNase (Sigma-Aldrich #D4527) and 100 mg/mL collagenase/dispase (Roche #10269638001) in PBS for 45 min at 37C. Cells were then sequentially filtered through 100- and 40-mm cell strainers (Falcon #352360 and #352340) and centrifuged at 1000 rpm for 5 min at 4C. Cells were resuspended in red blood cell lysis buffer (0.15 M NH4Cl, 10mM KHCO3, 0.1 mM EDTA) for 90 s at room temperature, followed by addition of DMEM (Gibco #11995-065) and FBS (Corning #MT35016CV). Cells were centrifuged at 1000 rpm for 5 min at 4C and then resuspended in PBS/10% FBS for further staining. The following antibodies were used: anti-CD31 APC, anti-CD45 APC, anti-Ly-6A/E (SCA1) APC/Cy7 (Thermo Fisher Scientific #BDB551262, #DB559864 and #560654), anti-CD326 (EpCAM) PE/Cy7 (Biolegend #118216) (all 1:100). DAPI (Sigma-Aldrich #D9542) was used to eliminate dead cells. Single stain controls and fluorescence-minus-one (FMO) controls were included for each experiment. For experiments that required the retention of a lung lobe for histology in addition to tissue processing for flow cytometry, the left lobe was isolated and removed using a hemostat (see below for Histology processing).

##### Lung tissue preparation, flow cytometry, FACS for Immune cells

For flow cytometry and sorting experiments of immune cell populations in the alveolar airspaces and the lungs, BALF (see below for method) recovery was performed and cells were placed on ice and either stained for BALF-only analysis or combined with lungs tissue digested as follows. Lungs were inflated by intratracheal instillation of 2 mL of Liberase buffer (0.1mg/ml Liberase Sigma Aldrich #05401127001, 50U/ml DNAse Sigma-Aldrich #D4527 in HBSS Sigma-Aldrich #05401127001). Lungs were placed on ice, minced, and incubated for 45 min at 37C. Cells were then sequentially filtered through 100- and 40-mm cell strainers (Falcon #352360 and #352340) and combined with the BALF-derived material. The complete samples were then centrifuged at 1000 rpm for 5 min at 4C. Cells were resuspended in red blood cell lysis buffer (0.15 M NH4Cl, 10mM KHCO3, 0.1 mM EDTA) for 90 s at room temperature, followed by addition of DMEM (Gibco #11995-065) and FBS (Corning #MT35016CV). Cells were centrifuged at 1000 rpm for 5 min at 4C and then resuspended in PBS/10% FBS for further staining. Samples were then blocked with anti-mouse CD16/32 antibody (BioLegend #101302) for 5 minutes on ice prior to cell surface staining. The following antibodies were used for the myeloid immune panel: anti-CD45 FITC (Thermo Fisher #553080) or anti-CD45 BV750 (BD #746947), anti-Cd11b BV605 (BD #563015), anti-CD64 APC (BioLegend #139306), anti-MerTK PE/Cy7(Fisher Scientific #25-5751-82), anti-SiglecF PE (BioLegend #15506), anti-Ly6G APC/Cy7 (BioLegend #127624) (all 1:100). DAPI (Sigma-Aldrich #D9542) was used to eliminate dead cells. Single stain controls and fluorescence-minus-one (FMO) controls were included for each experiment.

For detection of apoptotic cells, Annexin V-FITC Apoptosis Detection Kit (BD Biosciences #556547) was used according to the manufacturer’s instructions. After standard Epithelial staining, cells were washed with binding buffer and stained with Annexin V-FITC for 15 minutes at room temperature in the dark. Samples were then immediately analyzed by flow cytometry.

Flow cytometry analysis was performed on a BD LSR II or LSR Fortessa, and experiments that required sorting utilized a BD FACSAria II. Analysis of all samples was performed using BD FACSDiva and FlowJo software.

##### Bronchoalveolar lavage fluid (BALF) collection

Mice were euthanized by anesthetic overdose (avertin) and a 20G Angio-catheter (VWR #63019-889) was inserted through the trachea. BALF was obtained from animals by injecting and withdrawing 1 mL of warm 2% FBS, 2mM EDTA PBS solution (37°C), through the catheter, and repeated up to 5 times. BALF was then stored on ice until its use in flow cytometry, scRNA-seq, or MACS.

##### Macrophage depletion

Female C57/Bl6J mice were first anesthetized with isoflurane and sterile liposomes containing PBS or Clodronate (Liposoma BV #CP-005-005) were administered to animals via intratracheal administration at 50 uL per 20 g animal. Mice were given daily doses over 4 days. Naphthalene injury was performed 3 days after the last dose of liposomes, as described above.

##### Lung organoid co-cultures

AT2 cells were obtained via FACS (CD31-/CD45-/Ter119-/EpCAM+/Sca1-) from uninjured control or injured C57 Bl/6 and grown in co-culture conditions with stromal cells as previously described^26^. For co-culture conditions with alveolar macrophages, naïve DsRed mice were used as the source of AT2 cells (CD31-/CD45-/Ter119-/EpCAM+/Sca1-) while control or injured C57 Bl/6 mice were used as the source of AMs. AM co-cultures used media developed in the Tata lab^34^ and omitted IL-1β and EGF.

##### MACS enrichment of alveolar macrophages

BAL fluid was collected from uninjured control or injured animals as described and red blood cells were lysed (0.154M NH_4_Cl/10mM KHCO_3_/0.127mM EDTA). The remaining cell suspension was incubated with CD45 magnetic beads (Miltenyi #130-052-301) and positively selected using the OctoMACS (Miltenyi) platform. Cells were then passed through a 0.4μm cell strainer prior to lung organoid co-culture.

##### Histology and immunostaining

For histology of the lungs, mouse lung tissues were perfused, inflated with 0.8% low melting agarose (Thermo Scientific #16520050) in PBS, and fixed overnight at 4°C in 10% neutral-buffered formalin (Fisher Scientific #23-427098). Tissues were then switched to 70% ethanol prior to paraffin embedding. Formalin-fixed, paraffin embedded lung sections were cut to 5 µm thickness and mounted onto glass slides. Hematoxylin and eosin staining was performed where indicated. Sectioned lung tissues were stained for immunofluorescence as previously described^17^. Primary antibodies were incubated overnight at 4C at the indicated dilutions: rabbit anti-SPC (1:1000, Abcam #ab211362), mouse anti-acetyl-a-tubulin (1:100, Sigma-Aldrich #T7451), goat anti-CCSP (1:100, Santa Cruz Biotechnology #9772), rat anti-Ki67 (1:100 eBiosciences #14-5689-92), mouse anti-CCSP (1:100, Santa Cruz Biotechnology #390313), goat anti-GFP/YFP (1:400 AbCam #6673), rat anti-RAGE (1:100 R&D #MAB1179-SP), mouse anti-Cyp2f2 (1:100 Santa Cruz Biotechnology #374540), mouse anti-RFP/tdTOM (1:100, Invitrogen #MA5-15257). Slides were incubated with Alexa Fluor-coupled secondary antibodies for 1 h at room temperature (all Invitrogen, 1:200, see Key resources table). Slides were mounted using Prolong Gold with DAPI (Invitrogen # P36935). IF imaging was performed using a Nikon 90i Eclipse microscope with OptiScanIII motorized stage (Prior Scientific). Image analysis was done using ImageJ^52^.

##### Immunostaining of organoids in Matrigel

Transwell containing full sized organoids in Matrigel were fixed in 10% neutral buffered formalin (NBF) for at least 6 hours. Each transwell was then washed in PBS twice and submerged in Permeabilization Solution (PBS + 0.5% Triton) for one hour while gently rocking. Transwells were then washed once in Washing Solution (PBS + 0.2% Triton) and blocked in Blocking Solution (Washing Solution + 10% Normal Donkey Serum) at 4C while gently rocking (500ul on top and 500ul at the bottom). After overnight incubation, the Blocking Solution on top of each transwell was removed by decanting and substituted with Primary Antibody Solution (Washing Solution + Primary Antibodies). After 2 overnights at 4C, each transwell was then washed in Washing Solution and submerged in fresh Washing Solution for 30 minutes while rocking at room temperature. After 3 washes, 500ul of Blocking Solution was added at the bottom and 500ul of Secondary Antibody Solution (Washing Solution + Seconday Antibodies + DAPI) was added on top of each transwell. After an overnight incubation at 4C while rocking, each transwell was washed in Washing Solution and submerged in fresh Washing Solution for 30 minutes while rocking at room temperature. After 3 washes, the transwells were readied for imaging in PBS. The primary antibodies used were: rabbit anti-SPC (1:500, Abcam #ab211362) and rat anti-RAGE (1:50 R&D #MAB1179-SP). All secondary antibodies were Alexa Fluor-coupled secondary antibodies used at 1:100. Imaging was achieved by eliminating the excess buffer and flipping the transwells upside down using a Nikon 90i Eclipse microscope with OptiScanIII motorized stage (Prior Scientific). Image analysis was done using ImageJ^52^.

##### Generation and analysis of scRNA-seq data Mouse epithelial cell preparation

Female adult C57/Bl6J mice were injured with naphthalene or given Corn Oil control injections as described above and sacrificed at days 2 or 7 after injury. Epithelial populations were isolated from the CD31-/CD45-/EpCAM+/Sca1- and CD31-/CD45-/EpCAM+/Sca1+ gates by FACS (n = 2 mice per condition).

##### Mouse immune cell preparation

Female adult C57/Bl6J mice were injured with naphthalene or given control injections as described above and sacrificed at days 2 or 7 after injury. Before extracting lungs from the animals, BALF was obtained from the mice as described above and set aside on ice. The remaining lung tissue was dissociated and stained, and immune cells were isolated by FACS based on CD31-/CD119-/CD45+ surface markers. For sequencing, BALF was combined with sorted immune cells at a 1:1 ratio per animal (n=3 mice per condition).

##### Library preparation

ScRNA-seq was performed using the 10x Genomics platform. Cells were encapsulated with a 10X Genomics Chromium Controller Instrument and the following reverse transcription, cDNA amplification, and library preparation were performed using the Chromium Next GEM Single Cell 2’ v3 or v3.1 kits. Quantification and quality control of cDNA was determined using a QubitTM dsDNA HS assay kit and the Agilent TapeStation High Sensitivity D5000 ScreenTape System. Libraries were sequenced using an Illumina NextSeq500 at a depth of ∼50,000 reads/cell and later processed using CellRanger.

##### Analysis of scRNA-seq data

Analysis of raw CellRanger data was performed using Seurat v5^53^. Epithelial cells were filtered for UMI ≥ 1000, genes ≥ 500 and mitochondrial DNA ratio < 0.2. Immune cells were filtered for UMI ≥ 200, genes ≥ 100 and mitochondrial DNA ratio < 0.2. the Immune dataset was further filtered for doublets using scDblFinder. The remaining cells were re-clustered in their respective datasets and used for downstream analysis. For both datasets, differential gene expression analysis was performed with Seurat “FindMarkers” function to identify positively upregulated genes with a log fold change > 0.25.

##### Signatures Scores and Pathway Analysis

Proliferation score was calculated using “AddModuleScore” with the following genes: Mki67, Pcna, Top2a, Ccnb1, Ccnb2, Ccna2, Ccne1, Cdk1, Cdk2, Aurka, Aurkb, Cenpa, Cenpe, Cenpf, Ube2c, Bub1, Bub1b, Mcm2, Mcm3, Mcm4, Mcm5, Mcm6, Mcm7.

Primed AT2 score was calculated using “AddModuleScore” with the following genes: Lcn2, Il33, Lrg1, Dmkn, Ifi27l2a, Cxcl17, Itga7, Ly6e, Slpi, Atp5e.

Transitional AT2 score was calculated using “AddModuleScore” with the following genes: Krt8, Cldn4, Krt18, Krt19, Lgals3, Sfn, Anxa1, Myl12a, S100a6, Ybx1.

LAM (or foamy) score was calculated using “AddModuleScore” with the following genes: Trem2, Gpnmb, Spp1, Cd9, Fabp5, Apoe, Lpl, Lipa, Ctsb, Lgals3. The top genes from King et al, 2024 and Tleugabulova et al, 2024 were combined to obtain a 10-gene signature. Pathway analysis was performed using Reactome^39^.

##### Quantification and statistical analysis

The statistical tests adopted for each experiment are listed in the legend of each figure. Data are represented as mean ± SEM unless otherwise noted. The following notations are used to indicate statistical significance: * = p-value < 0.05, ** = p-value < 0.01, *** = p-value < 0.001 and **** = p-value < 0.0001. Absence of asterisk means no significance compared to control.

---

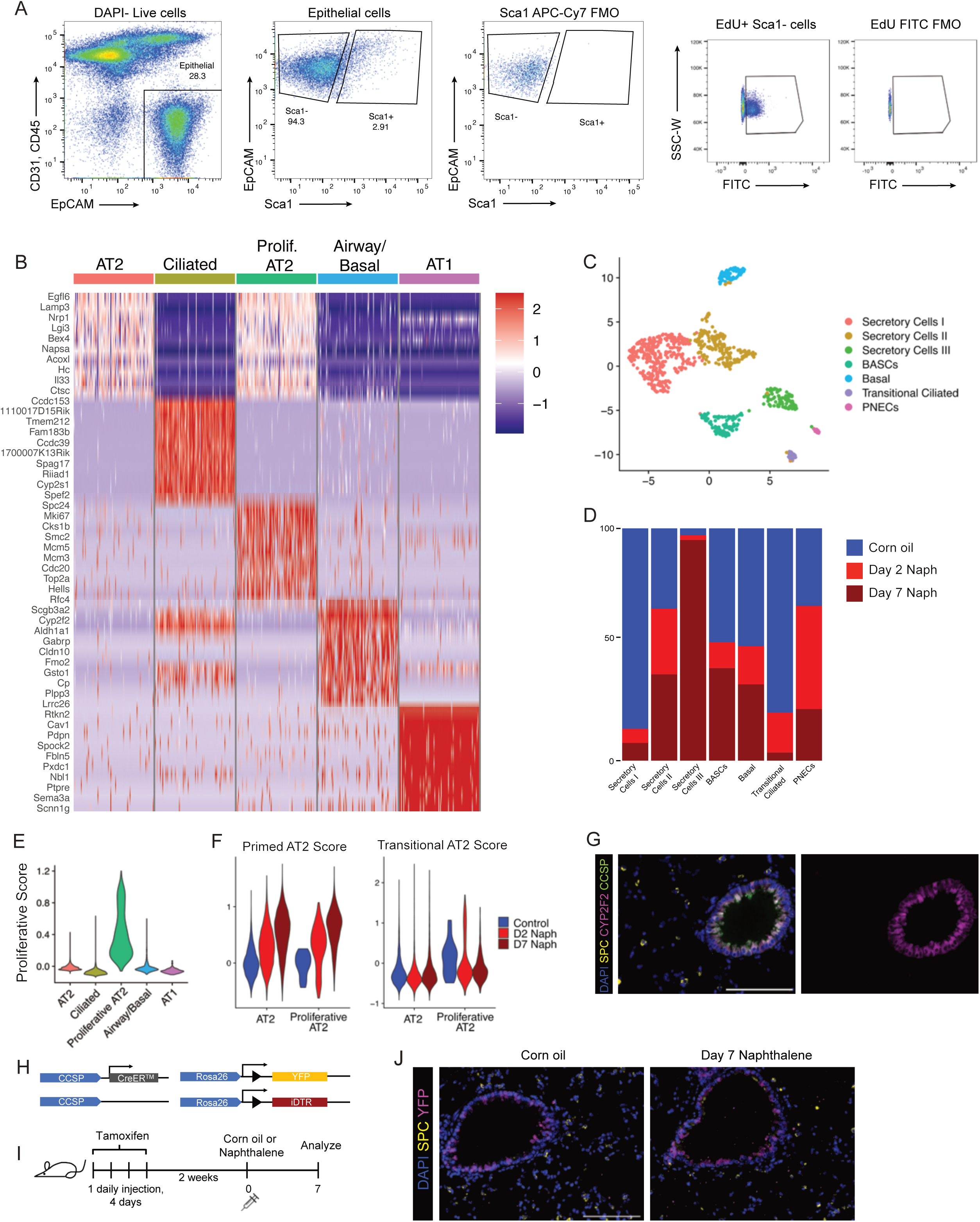

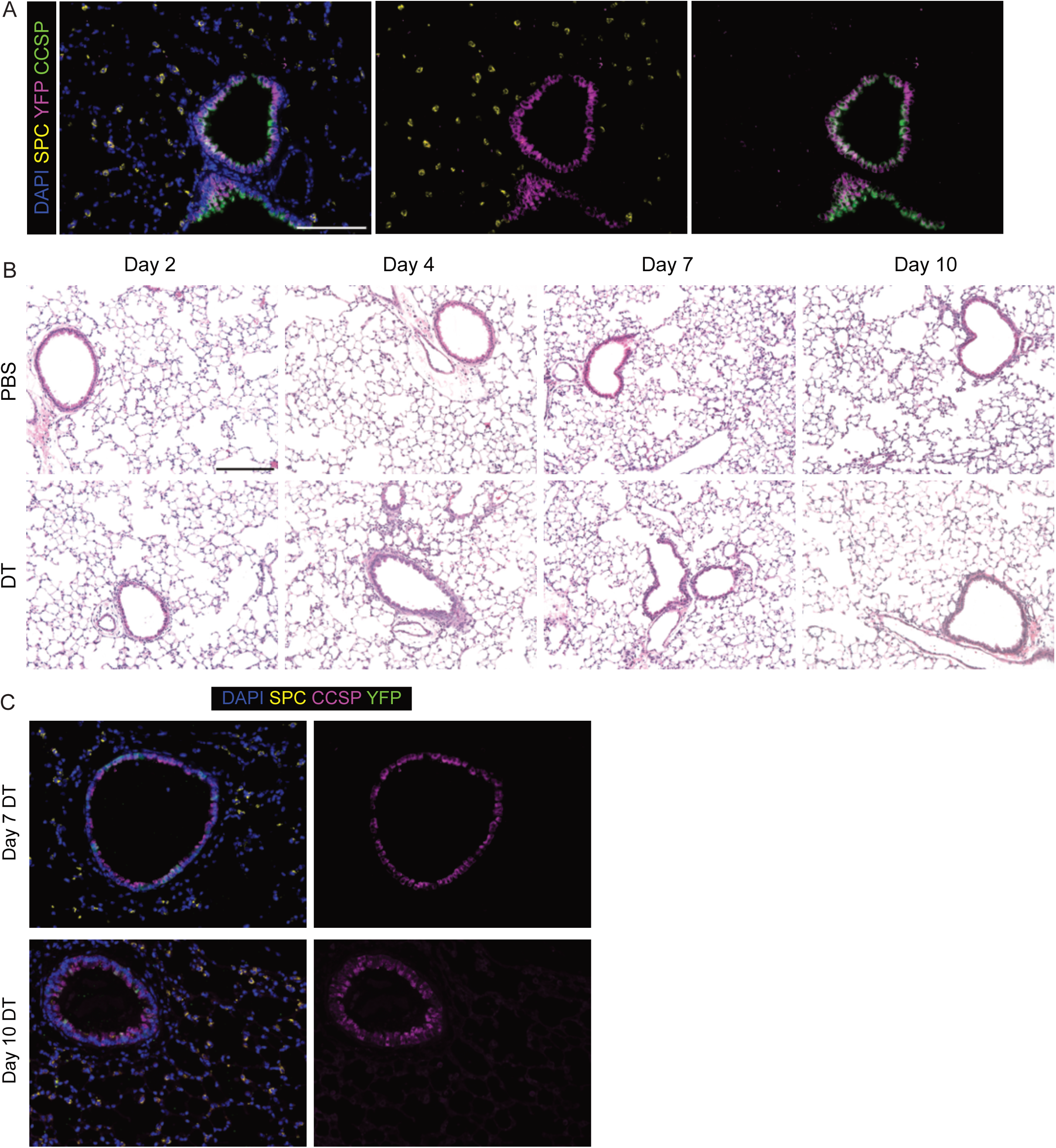

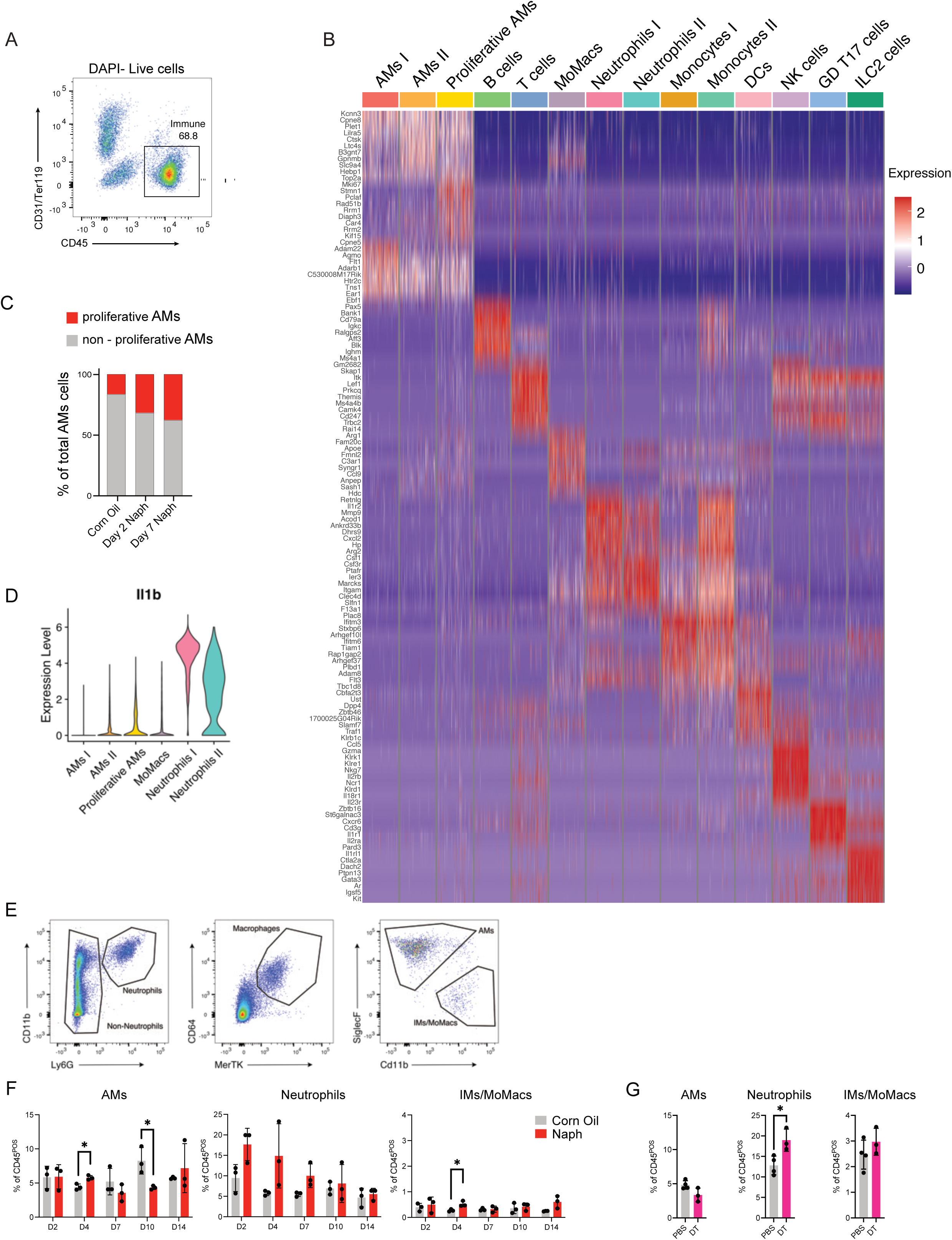

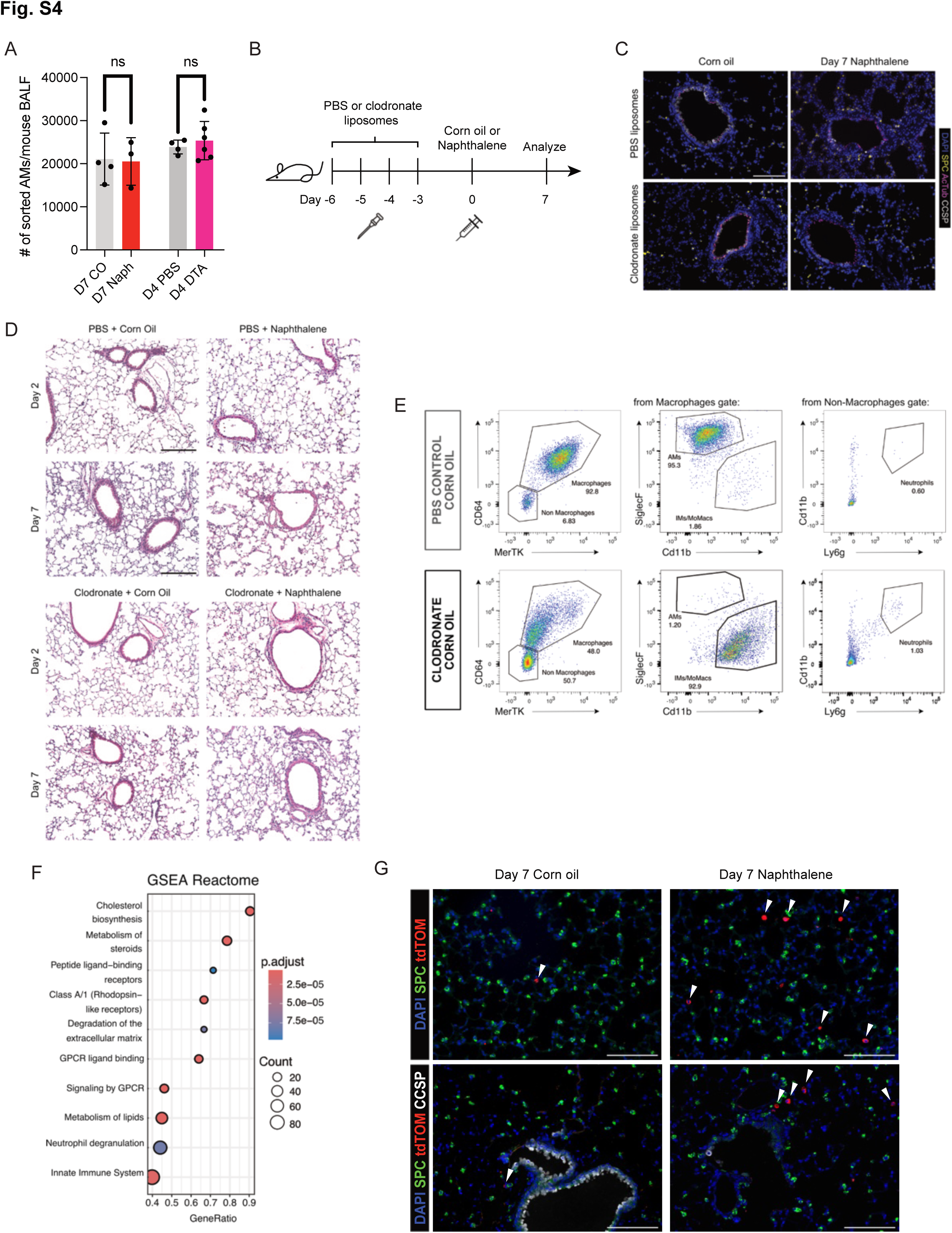

## References

1. Butler, J.P., Loring, S.H., Patz, S., Tsuda, A., Yablonskiy, D.A., and Mentzer, S.J. (2012). Evidence for adult lung growth in humans. N. Engl. J. Med. 367, 244–247.

2. Mentzer, S.J. (2018). The puzzling mechanism of compensatory lung growth. Stem Cell Investig 5, 8.

3. Sun, F., and Poss, K.D. (2023). Inter-organ communication during tissue regeneration. Development 150. 10.1242/dev.202166.

4. Rodgers, J.T., King, K.Y., Brett, J.O., Cromie, M.J., Charville, G.W., Maguire, K.K., Brunson, C., Mastey, N., Liu, L., Tsai, C.-R., et al. (2014). mTORC1 controls the adaptive transition of quiescent stem cells from G0 to G(Alert). Nature 510, 393–396.

5. Johnson, K., Bateman, J., DiTommaso, T., Wong, A.Y., and Whited, J.L. (2018). Systemic cell cycle activation is induced following complex tissue injury in axolotl. Dev. Biol. 433, 461–472.

6. Sikkema, L., Ramírez-Suástegui, C., Strobl, D.C., Gillett, T.E., Zappia, L., Madissoon, E., Markov, N.S., Zaragosi, L.-E., Ji, Y., Ansari, M., et al. (2023). An integrated cell atlas of the lung in health and disease. Nat. Med., 1–15.

7. Barkauskas, C.E., Cronce, M.J., Rackley, C.R., Bowie, E.J., Keene, D.R., Stripp, B.R., Randell, S.H., Noble, P.W., and Hogan, B.L.M. (2013). Type 2 alveolar cells are stem cells in adult lung. J. Clin. Invest. 123, 3025–3036.

8. Rawlins, E.L., Ostrowski, L.E., Randell, S.H., and Hogan, B.L.M. (2007). Lung development and repair: contribution of the ciliated lineage. Proc. Natl. Acad. Sci. U. S. A. 104, 410–417.

9. Whitsett, J.A. (2018). Airway Epithelial Differentiation and Mucociliary Clearance. Ann. Am. Thorac. Soc. 15, S143–S148.

10. Guha, A., Deshpande, A., Jain, A., Sebastiani, P., and Cardoso, W.V. (2017). Uroplakin 3a+ Cells Are a Distinctive Population of Epithelial Progenitors that Contribute to Airway Maintenance and Post-injury Repair. Cell Rep. 19, 246–254.

11. Basil, M.C., Katzen, J., Engler, A.E., Guo, M., Herriges, M.J., Kathiriya, J.J., Windmueller, R., Ysasi, A.B., Zacharias, W.J., Chapman, H.A., et al. (2020). The Cellular and Physiological Basis for Lung Repair and Regeneration: Past, Present, and Future. Cell Stem Cell 26, 482–502.

12. Lechner, A.J., Driver, I.H., Lee, J., Conroy, C.M., Nagle, A., Locksley, R.M., and Rock, J.R. (2017). Recruited Monocytes and Type 2 Immunity Promote Lung Regeneration following Pneumonectomy. Cell Stem Cell 21, 120–134.e7.

13. Engler, A.E., Ysasi, A.B., Pihl, R.M.F., Villacorta-Martin, C., Heston, H.M., Richardson, H.M.K., Thapa, B.R., Moniz, N.R., Belkina, A.C., Mazzilli, S.A., et al. (2020). Airway-Associated Macrophages in Homeostasis and Repair. Cell Rep. 33, 108553.

14. Dagher, R., Copenhaver, A.M., Besnard, V., Berlin, A., Hamidi, F., Maret, M., Wang, J., Qu, X., Shrestha, Y., Wu, J., et al. (2020). IL-33-ST2 axis regulates myeloid cell differentiation and activation enabling effective club cell regeneration. Nat. Commun. 11, 4786.

15. Reyes, N.S., Krasilnikov, M., Allen, N.C., Lee, J.Y., Hyams, B., Zhou, M., Ravishankar, S., Cassandras, M., Wang, C., Khan, I., et al. (2022). Sentinel *p16*^INK4a+^ cells in the basement membrane form a reparative niche in the lung. Science 378, 192–201.

16. Lucas, C.D., Medina, C.B., Bruton, F.A., Dorward, D.A., Raymond, M.H., Tufan, T., Etchegaray, J.I., Barron, B., Oremek, M.E.M., Arandjelovic, S., et al. (2022). Pannexin 1 drives efficient epithelial repair after tissue injury. Sci Immunol 7, eabm4032.

17. Louie, S.M., Moye, A.L., Wong, I.G., Lu, E., Shehaj, A., Garcia-de-Alba, C., Ararat, E., Raby, B.A., Lu, B., Paschini, M., et al. (2022). Progenitor potential of lung epithelial organoid cells in a transplantation model. Cell Rep. 39, 110662.

18. Choi, J., Park, J.-E., Tsagkogeorga, G., Yanagita, M., Koo, B.-K., Han, N., and Lee, J.-H. (2020). Inflammatory signals induce AT2 cell-derived damage-associated transient progenitors that mediate alveolar regeneration. Cell Stem Cell 27, 366–382.e7.

19. Strunz, M., Simon, L.M., Ansari, M., Kathiriya, J.J., Angelidis, I., Mayr, C.H., Tsidiridis, G., Lange, M., Mattner, L.F., Yee, M., et al. (2020). Alveolar regeneration through a Krt8+ transitional stem cell state that persists in human lung fibrosis. Nat. Commun. 11, 3559.

20. Kobayashi, Y., Tata, A., Konkimalla, A., Katsura, H., Lee, R.F., Ou, J., Banovich, N.E., Kropski, J.A., and Tata, P.R. (2020). Persistence of a regeneration-associated, transitional alveolar epithelial cell state in pulmonary fibrosis. Nat. Cell Biol. 22, 934–946.

21. Buckpitt, A., Boland, B., Isbell, M., Morin, D., Shultz, M., Baldwin, R., Chan, K., Karlsson, A., Lin, C., Taff, A., et al. (2002). Naphthalene-induced respiratory tract toxicity: metabolic mechanisms of toxicity. Drug Metab. Rev. 34, 791–820.

22. Buckpitt, A., Kephalopoulos, S., Koistinen, K., Kotzias, D., Morawska, L., and Sagunski, H. (2010). Naphthalene (World Health Organization).

23. Gerdes, J., Lemke, H., Baisch, H., Wacker, H.H., Schwab, U., and Stein, H. (1984). Cell cycle analysis of a cell proliferation-associated human nuclear antigen defined by the monoclonal antibody Ki-67. J. Immunol. 133, 1710–1715.

24. Rawlins, E.L., Okubo, T., Xue, Y., Brass, D.M., Auten, R.L., Hasegawa, H., Wang, F., and Hogan, B.L.M. (2009). The Role of Scgb1a1+ Clara Cells in the Long-Term Maintenance and Repair of Lung Airway, but Not Alveolar, Epithelium. Cell Stem Cell 4, 525–534.

25. Buch, T., Heppner, F.L., Tertilt, C., Heinen, T.J.A.J., Kremer, M., Wunderlich, F.T., Jung, S., and Waisman, A. (2005). A Cre-inducible diphtheria toxin receptor mediates cell lineage ablation after toxin administration. Nat. Methods 2, 419–426.

26. Lee, J.-H., Bhang, D.H., Beede, A., Huang, T.L., Stripp, B.R., Bloch, K.D., Wagers, A.J., Tseng, Y.-H., Ryeom, S., and Kim, C.F. (2014). Lung Stem Cell Differentiation in Mice Directed by Endothelial Cells via a BMP4-NFATc1-Thrombospondin-1 Axis. Cell 156, 440–455.

27. Hoagland, D.A., Rodríguez-Morales, P., Mann, A.O., Baez Vazquez, A.Y., Yu, S., Lai, A., Kane, H., Dang, S.M., Lin, Y., Thorens, L., et al. (2025). Macrophage-derived oncostatin M repairs the lung epithelial barrier during inflammatory damage. Science 389, 169–175.

28. Hung, L.-Y., Sen, D., Oniskey, T.K., Katzen, J., Cohen, N.A., Vaughan, A.E., Nieves, W., Urisman, A., Beers, M.F., Krummel, M.F., et al. (2019). Macrophages promote epithelial proliferation following infectious and non-infectious lung injury through a Trefoil factor 2-dependent mechanism. Mucosal Immunol. 12, 64–76.

29. Rodríguez-Morales, P., and Franklin, R.A. (2023). Macrophage phenotypes and functions: resolving inflammation and restoring homeostasis. Trends Immunol. 44, 986–998.

30. Lee, J.S., Koh, J.-Y., Yi, K., Kim, Y.-I., Park, S.-J., Kim, E.-H., Kim, S.-M., Park, S.H., Ju, Y.S., Choi, Y.K., et al. (2021). Single-cell transcriptome of bronchoalveolar lavage fluid reveals sequential change of macrophages during SARS-CoV-2 infection in ferrets. Nat. Commun. 12, 4567.

31. Wang, L., Netto, K.G., Zhou, L., Liu, X., Wang, M., Zhang, G., Foster, P.S., Li, F., and Yang, M. (2021). Single-cell transcriptomic analysis reveals the immune landscape of lung in steroid-resistant asthma exacerbation. Proc. Natl. Acad. Sci. U. S. A. 118, e2005590118.

32. Aegerter, H., Lambrecht, B.N., and Jakubzick, C.V. (2022). Biology of lung macrophages in health and disease. Immunity 55, 1564–1580.

33. Van Hoecke, L., Job, E.R., Saelens, X., and Roose, K. (2017). Bronchoalveolar Lavage of Murine lungs to analyze inflammatory cell infiltration. J. Vis. Exp. 10.3791/55398.

34. Konishi, S., Tata, A., and Tata, P.R. (2022). Defined conditions for long-term expansion of murine and human alveolar epithelial stem cells in three-dimensional cultures. STAR Protoc. 3, 101447.

35. Leslie, J., Millar, B.J., Del Carpio Pons, A., Burgoyne, R.A., Frost, J.D., Barksby, B.S., Luli, S., Scott, J., Simpson, A.J., Gauldie, J., et al. (2020). FPR-1 is an important regulator of neutrophil recruitment and a tissue-specific driver of pulmonary fibrosis. JCI Insight 5. 10.1172/jci.insight.125937.

36. Misharin, A.V., Morales-Nebreda, L., Reyfman, P.A., Cuda, C.M., Walter, J.M., McQuattie-Pimentel, A.C., Chen, C.-I., Anekalla, K.R., Joshi, N., Williams, K.J.N., et al. (2017). Monocyte-derived alveolar macrophages drive lung fibrosis and persist in the lung over the life span. J. Exp. Med. 214, 2387–2404.

37. Thepen, T., Van Rooijen, N., and Kraal, G. (1989). Alveolar macrophage elimination in vivo is associated with an increase in pulmonary immune response in mice. J. Exp. Med. 170, 499–509.

38. Van Rooijen, N., and Sanders, A. (1994). Liposome mediated depletion of macrophages: mechanism of action, preparation of liposomes and applications. J. Immunol. Methods 174, 83–93.

39. Milacic, M., Beavers, D., Conley, P., Gong, C., Gillespie, M., Griss, J., Haw, R., Jassal, B., Matthews, L., May, B., et al. (2024). The reactome pathway knowledgebase 2024. Nucleic Acids Res. 52, D672–D678.

40. King, E.M., Zhao, Y., Moore, C.M., Steinhart, B., Anderson, K.C., Vestal, B., Moore, P.K., McManus, S.A., Evans, C.M., Mould, K.J., et al. (2024). Gpnmb and Spp1 mark a conserved macrophage injury response masking fibrosis-specific programming in the lung. JCI Insight 9. 10.1172/jci.insight.182700.

41. Cui, H., Jiang, D., Banerjee, S., Xie, N., Kulkarni, T., Liu, R.-M., Duncan, S.R., and Liu, G. (2020). Monocyte-derived alveolar macrophage apolipoprotein E participates in pulmonary fibrosis resolution. JCI Insight 5. 10.1172/jci.insight.134539.

42. Wohnhaas, C.T., Baßler, K., Watson, C.K., Shen, Y., Leparc, G.G., Tilp, C., Heinemann, F., Kind, D., Stierstorfer, B., Delić, D., et al. (2024). Monocyte-derived alveolar macrophages are key drivers of smoke-induced lung inflammation and tissue remodeling. Front. Immunol. 15, 1325090.

43. Cruz Tleugabulova, M., Melo, S.P., Wong, A., Arlantico, A., Liu, M., Webster, J.D., Lau, J., Lechner, A., Corak, B., Hodgins, J.J., et al. (2024). Induction of a distinct macrophage population and protection from lung injury and fibrosis by Notch2 blockade. Nat. Commun. 15, 9575.

44. Español-Suñer, R., Carpentier, R., Van Hul, N., Legry, V., Achouri, Y., Cordi, S., Jacquemin, P., Lemaigre, F., and Leclercq, I.A. (2012). Liver progenitor cells yield functional hepatocytes in response to chronic liver injury in mice. Gastroenterology 143, 1564–1575.e7.

45. Pervizaj-Oruqaj, L., Selvakumar, B., Ferrero, M.R., Heiner, M., Malainou, C., Glaser, R.D., Wilhelm, J., Bartkuhn, M., Weiss, A., Alexopoulos, I., et al. (2024). Alveolar macrophage-expressed Plet1 is a driver of lung epithelial repair after viral pneumonia. Nat. Commun. 15, 87.

46. Meiners, S., Eickelberg, O., and Königshoff, M. (2015). Hallmarks of the ageing lung. Eur. Respir. J. 45, 807–827.

47. Kim, C.F.B., Jackson, E.L., Woolfenden, A.E., Lawrence, S., Babar, I., Vogel, S., Crowley, D., Bronson, R.T., and Jacks, T. (2005). Identification of Bronchioalveolar Stem Cells in Normal Lung and Lung Cancer. Cell 121, 823–835.

48. Love, M.E., and Proud, D. (2022). Respiratory viral and bacterial exacerbations of COPD-the role of the airway epithelium. Cells 11, 1416.

49. Ghosh, M.C., Gorantla, V., Makena, P.S., Luellen, C., Sinclair, S.E., Schwingshackl, A., and Waters, C.M. (2013). Insulin-like growth factor-I stimulates differentiation of ATII cells to ATI-like cells through activation of Wnt5a. Am. J. Physiol. Lung Cell. Mol. Physiol. 305, L222–8.

50. Chung, E.J., Kwon, S., Reedy, J.L., White, A.O., Song, J.S., Hwang, I., Chung, J.Y., Ylaya, K., Hewitt, S.M., and Citrin, D.E. (2021). IGF-1 receptor signaling regulates type II pneumocyte senescence and resulting macrophage polarization in lung fibrosis. Int. J. Radiat. Oncol. Biol. Phys. 110, 526–538.

51. Gao, F., Li, C., Smith, S.M., Peinado, N., Kohbodi, G., Tran, E., Loh, Y.-H.E., Li, W., Borok, Z., and Minoo, P. (2022). Decoding the IGF1 signaling gene regulatory network behind alveologenesis from a mouse model of bronchopulmonary dysplasia. Elife 11. 10.7554/eLife.77522.

52. Schneider, C.A., Rasband, W.S., and Eliceiri, K.W. (2012). NIH Image to ImageJ: 25 years of image analysis. Nat. Methods 9, 671–675.

53. Hao, Y., Stuart, T., Kowalski, M.H., Choudhary, S., Hoffman, P., Hartman, A., Srivastava, A., Molla, G., Madad, S., Fernandez-Granda, C., et al. (2024). Dictionary learning for integrative, multimodal and scalable single-cell analysis. Nat. Biotechnol. 42, 293–304.

