## Supplemental Figures Legends for "Airway injury induces alveolar epithelial responses mediated by macrophages"

**Supplementary Information**

**Figure S1, related to Figure 1: Transient proliferation of AT2 cells after naphthalene airway injury.**

A. Scatter plot of FACS gating strategy for Sca1- and Sca1+ lung epithelial cells and Edu^POS^ gate for proliferating cells.
B. Heatmap of the top 10 markers for each cell cluster of merged lung epithelium scRNA-seq as shown in Figure 1C (100 randomly picked cells are shown per each cluster).
C. UMAP plot and annotation of re-clustered Airway/Basal cluster from 1C.
D. Composition of clusters shown in S1C by condition.
E. Proliferation score by cluster of lung epithelium scRNA-seq shown in Figure 1C.
F. Violin Plots of Primed AT2 and transitional AT2 score signatures for the AT2 clusters from Figure 1C.
G. IF images of normal mouse lungs. AW = airway. Scale bar = 100 µm.
H. Diagram of alleles of the CCSP-CreER+/-; Rosa26-LSL-YFP/iDTR mice. Triangle represents LoxP-Stop-LoxP sites.
I. Schematic of lineage tracing club cells after corn oil control or naphthalene injury in CCSP-CreER/-; Rosa26-LSL-YFP/iDTR mice.
J. IF images of lineage-traced club cells after corn oil control or naphthalene injury as described in (F). Scale bar = 100 µm.

**Figure S2, related to Figure 2: Transient proliferation of AT2 cells after genetic club cell ablation.**

A. IF of CCSP-CreER/-; Rosa26-LSL-YFP/iDTR mouse lungs after tamoxifen induction at the day 0 timepoint indicated in Figure 2A. Mice were not administered PBS or DT. Scale bar = 100 µm.
B. H&E staining of mouse lungs after genetic club cell ablation at indicated timepoints. Scale bar = 200 µm.
C. IF images of mouse lungs at the indicated timepoints after genetic club cell ablation. Scale bar = 100 µm.

**Figure S3, related to Figure 3: Multiple myeloid lineages respond to airway injury.**

A. FACS strategy used to isolate CD45+ immune cells from the lungs of mice treated with corn oil control or naphthalene.
B. Heatmap of the top 10 markers for each cell cluster of merged lung BALF fluid cells and whole lung immune cell scRNA-seq as shown in Figure 3B.
C. Percentage of proliferative cells of the total number of AMs for each condition according to scRNA-seq data. Grey: non proliferative AMs fraction, Red: proliferative AM fraction.
D. Violin Plot for the expression of Il1B across the Myeloid populations from Figure 1B.
E. Gating strategy for identification of AMs, Neutrophils, IMs/MoMacs in whole lung digestion after naphthalene injury.
F. Quantification of AMs, Neutrophils and IMs/MoMacs in whole lung digestion at multiple time points after naphthalene injury.
G. Quantification of AMs, Neutrophils and IMs/MoMacs in whole lung digestion 2 days after induction and injury of CCSP-CreER+/-; Rosa26-LSL-YFP/iDTR mice

**Figure S4, related to Figure 4: Cell populations in the airspace mediate AT2 cell proliferation after airway injury.**

A. Comparison of the number of sorted AMs from indicated conditions. N = 3-6 mice per condition. Student’s T-test: ns = no significance
B. Schematic of liposome administration in mice prior to control or naphthalene injury. PBS control or clodronate liposome was administered daily via intratracheal injection.
C. IF images of mice that received liposome pre-treatment prior to control or naphthalene injury, sacrificed at day 7 post-injury. Scale bar = 100 µm.
D. Hematoxylin and eosin images of mice that received liposome pre-treatment prior to control or naphthalene injury. Scale bar = 200 µm.
E. Representative flow cytometry analysis of Myeloid populations from the BALF of mice that received PBS or liposome pre-treatment prior to Corn Oil control, sacrificed at day 7 post-injury.
F. GSEA Reactome pathways enriched in AMs II vs AMs I cluster.
G. IF images of Spp1-CreERT2; LSL-tdTOM sacrificed 7 days post Naphthalene injury and treated with tamoxifen at -1, 0, +1 and +4 days from injury. Top panels: alveolar space. Bottom panel: airway regions.

**Table S1, related to Figure 1: Differentially expressed genes in Epithelium scRNAseq.**

Results of differential gene expression analysis of epithelial cell clusters in Figure 1C.

**Table S2, related to Figure 1: Differentially expressed genes after re-clustering of the Airway/Basal cells from Figure 1C.**

Results of differential gene expression analysis of epithelial cell clusters in Figure S1C.

**Table S3, related to Figure 3: Differentially expressed genes in Immune scRNAseq.**

Results of differential gene expression analysis of immune cell clusters in Figure 3B.

**Table S4, related to Figure 3: Differentially expressed genes in AMs I vs AM II.**

**Table S5, related to Figure 3: Differentially expressed genes in Neutrophils I vs Neutrophils II.**
